# Opportunities and challenges in using remote sensing for identifying grassland restoration sites and invasive tree species management in a global biodiversity hotspot

**DOI:** 10.1101/2020.07.24.219535

**Authors:** M. Arasumani, Milind Bunyan, V. V. Robin

## Abstract

Tropical montane grasslands (TMG) support biodiverse and endemic taxa and provide vital ecosystems services to downstream communities. Yet invasive alien tree species across the world have threatened tropical grasslands and grassland endemic species. In India, TMG in the Shola Sky Islands of the Western Ghats have been reduced due to exotic tree invasions (acacias, pines, and eucalyptus species). The loss of grassland habitat has, in turn, reduced the range sizes of species endemic to grasslands (plants, birds, amphibians, and mammals), driving some populations to local extinction. Grassland conversion to exotic trees has also reduced the annual runoff in the Western Ghats. Conserving existing grassland and restoring invaded habitat is critical to reverse these losses. This research focused on identifying grassland restoration sites using satellite images with a high spatial resolution (RapidEye). We used an object-oriented Random Forest classification to map the area for grassland restoration. We have identified an area of 254 sq. km. as suitable for grassland restoration and an area of 362 sq. km. for grassland conservation and prevention from invasion by exotic tree species. For restoration, we recommend a cautious removal of young and isolated exotic trees at the invasion front and restoring grasslands, instead of removing dense stands of mature exotic trees. We find that areas with low fire frequency areas tend to be invaded, but areas invaded by exotic trees tend to burn hotter which may be harmful to grassland species and ecosystems. We assume that removing exotic tree species in the identified restoration sites and restoring the grassland will be helpful in recovering lost habitat and ensuring the viability of populations of indigenous and endemic species and increasing streamflow.

## 1. Introduction

Tropical montane grasslands (TMG) are high elevation grasslands forming only 2% of all the grasslands in the world (Dixon et al., 2014) and can be found in Brazil (de Abreu and Durigan, 2011; Koch et al., 2016), Colombia (Farley, 2007), Australia (Fairfax et al., 2009), the Eastern Arc Mountains (Pellikka et al., 2009), the Hawaiian Islands (Daehler, 2005) and India (Arasumani et al., 2019). TMG support high endemism (Brooks et al., 2006), regulate the global carbon cycle (Gibson, 2009) and serve as a source of water to downstream communities (Robin and Nandini, 2012). Yet TMG have also been altered by human activities for decades (Arasumani et al., 2019; Hermann et al., 2016). Despite this, these grasslands do not benefit from the conservation and restoration efforts afforded to tropical montane forests, possibly due to the limited information on these grasslands (Joshi et al., 2018). In India, TMG have even been classified as wastelands in forest management plans as they are unlikely to generate revenue, contrary to the timber (even if exotic) found in the forests (Joshi et al., 2018).

### 1.1. Tropical montane grasslands loss due to invasive exotic trees

In recent times TMG face a novel threat through the establishment and expansion of exotic tree plantations (Arasumani et al., 2019; Ngorima and Shackleton, 2019). Exotic trees that were planted across TMG (Richardson and Van Wilgen, 2004) have now become invasive in tropical grasslands (Arasumani et al., 2019) aggravating grassland loss. Invasive alien species (IAS) threaten biodiversity and human livelihoods globally (Early et al., 2016; Shackleton et al., 2019; Verbrugge et al., 2019) and have been shown to significantly modify ecosystem structure and function by altering nutrient cycles and vegetation patterns (Richardson and Bond, 1991; Richardson et al., 1994). In many tropical and subtropical regions, acacias and pines have invaded large swathes of grassland including South Africa, India, Colombia, Brazil and Australia (Arasumani et al., 2019; Gwate et al., 2016; Richardson, 1998). For instance in South Africa, more than half of the montane grasslands were lost to exotic trees over a 60-year period (Weyer et al., 2015). In Brazil, 17% of the montane grasslands were lost to tree plantations in the Campos de Cima da Serra (Hermann et al., 2016). Similarly, in the Western Ghats in India, 23% of the montane grasslands were converted into invasive exotic trees over 44 years (Arasumani et al., 2019).

Invasion by exotic trees not only reduce the extent of grassland but also threaten endemic grassland species (Allan et al., 1997; Robin et al., 2014), alter hydrological regimes (Buytaert et al., 2007; Le Maitre et al., 2000; Sikka et al., 2003), reduce wildlife grazing capacity (Yapi et al., 2018) and impact livelihood practices of traditional communities (Cordero et al., 2018; Shackleton et al., 2019). Exotic tree invasions have also been reported to increase fuel loads and fire intensities (Van Wilgen and Richardson, 1985), which degrade soil quality (Lazzaro et al., 2014) and lead to soil erosion (Van der Waal et al., 2012).

### 1.2. Restoration of montane grasslands from invasive exotic trees

It is imperative to conserve and restore the last-remaining grasslands while prioritizing the locations of these efforts. The first step is to assess the status to which the grassland has been modified by exotic tree invasion (Le Maitre et al., 2011). Efficient grassland restoration also needs an understanding of the dynamics and drivers that have caused ecosystem modification and landscape change (Arasumani et al., 2019). To date, attempts to restore montane grasslands from exotic tree invasions have incorporated approaches that are both passive (i.e. preventing the introduction and spread of non-native species (Cuevas and Zalba, 2010)); and active (i.e. biological control (Le Maitre et al., 2011)). Most restoration plans are however passive, and attempt to restore invaded grasslands by eliminating existing invaders and preventing their regeneration (Le Maitre et al., 2011).

For invasive species such as *Acacia mearnsii* that grow rapidly and disperse seeds widely, removing mature trees is often ineffective in restoring invaded grasslands (Le Maitre et al., 2011). An approach that targets the removal of young (small) exotic trees—the invasion front—from the grasslands would be more effective at restoring grasslands instead (Souza-Alonso et al., 2017). Similarly restoring grasslands where isolated, but mature trees exist in grassland patches could be an easy way to restrict further dispersal from these trees.

### 1.3. Montane grasslands in the Shola Sky Islands of the Western Ghats

The Shola Sky Islands are a mosaic of montane grasslands and forests (Robin and Nandini, 2012) and support highly endemic and endangered species (Robin and Nandini, 2012). Most of the major southern Indian rivers also originate from the shola grasslands (Robin and Nandini, 2012), benefitting millions of downstream users. The montane grasslands of the Western Ghats contain at least 70 threatened grass species (Matthew, 1999) with 30 of these listed as endangered (Karunakaran et al., 1998). The shola grassland also supports an endangered ungulate (Nilgiri tahr, *Nilgiritragus hylocrius*; (Alempath, 2008)) and a threatened bird (Nilgiri Pipit, *Anthus nilghiriensis*; (Lele et al.; Robin et al., 2014)). Additionally, there are at least 20 frog species that are restricted to these grasslands (Abraham et al., 2015; Biju et al., 2010; Princy et al., 2017; Vijayakumar et al., 2014). Although locally common, all these species are specialists that are restricted to the narrow ecological range of the montane grasslands of the Shola Sky Islands.

Over the last few decades, however, 340 sq. km. of montane grasslands in the Western Ghats were lost to agricultural expansion, exotic tree plantations and the subsequent invasion from these plantations (Arasumani et al., 2018; Arasumani et al., 2019; Joshi et al., 2018), impacting associated taxa (Alempath, 2008; Robin et al., 2014). Currently, a couple of state forest departments are planning major restoration initiatives in these habitats and there is much public concern over this landscape change, including a state high court public interest litigation, driven by concerns of water security and biodiversity loss. State land management agencies, however, may not have sufficient information on identifying suitable sites for restoration. Large parts of the landscape are difficult to access and different sites may even require different tools and strategies (e.g. machine-based or manual uprooting), which will be determined by the stage of invasion by these exotic trees. Developing a reasonably accurate estimate of the restoration effort (including finances) is essential to improving the odds of successful restoration. There are however, two major challenges with developing an appropriate restoration effort. The first is identifying potential restoration sites at landscape scales; this would require an approach using remotely sensed data. The second, which is equally critical, is monitoring and limiting the spread of invasion while restoration takes place.

In order to use remote sensing data to overcome the first challenge, it is essential to detect specific features associated with the landscape in an image. Traditional supervised and unsupervised classifications of remote sensing images were developed for a pixel-based classification (PBC) approach, but these methods are often not suitable for classifying high-resolution satellite images due to absence of spatial shape and texture information (Blaschke and Strobl, 2001). A PBC is prone to salt-and-pepper error with high resolution images as single (or very small groups of) pixels are erroneously classified into different categories.

An Object-Oriented Classification (OOC) classifies objects into homogeneous regions, reducing such salt-and-pepper errors (Niphadkar et al., 2017) and numerous studies suggest that OOC is better than PBC for classifying high-resolution images (Blaschke and Strobl, 2001). Most of these studies have however focused on classifying discrete landscape units with regular shapes or homogeneous regions such as urban habitats, forest types, and water bodies. We are unaware of studies that have used remote sensing techniques for identifying grassland restoration sites in a complex mixed environment. In this study we endeavour to assess the viability of this approach at landscape scales.

Tackling the second challenge (i.e. preventing subsequent invasion) on the other hand, requires identifying the parameters that influence the spread of invasive plants (i.e. the invasion front), into the grasslands. This information is critical to prevent the continuing spread of invasives while restoration activities are ongoing. Previous landscape-level analyses with satellite image-based classified data have indicated various different factors to be responsible for the presence of invasives, including distance to source trees, topography and fire history (Arasumani et al., 2018; Van Wilgen and Richardson, 1985). We investigate the influence of these factors on the invasion front.

In this study therefore, we aim to identify sites for grassland restoration and identify variables associated with the invasion front of exotic invasive trees. Specifically, we examine the utility of satellite imagery to a) identify the invasion front and isolated mature trees in grasslands as restoration sites and b) landscape features associated with the current invasion front.

## 2. Methods

### 2.1. Study Area

A previous study (Arasumani et al., 2019) had examined exotic tree planting and invasion across the Shola Sky Islands and found that it was restricted to the two largest island-complexes the Nilgiris and the Palani - Annamalai Hills. For this study, we focused on these two complexes (Fig.1), with a study area covering approximately 2923 sq. km. of montane ecosystems of the Western Ghats, above 1400 amsl of the Western Ghats (Robin and Nandini, 2012). We delineated the study area (above 1400 amsl) by reclassifying the ASTER GDEM images in ArcGIS 10.5.

**Fig. 1.**
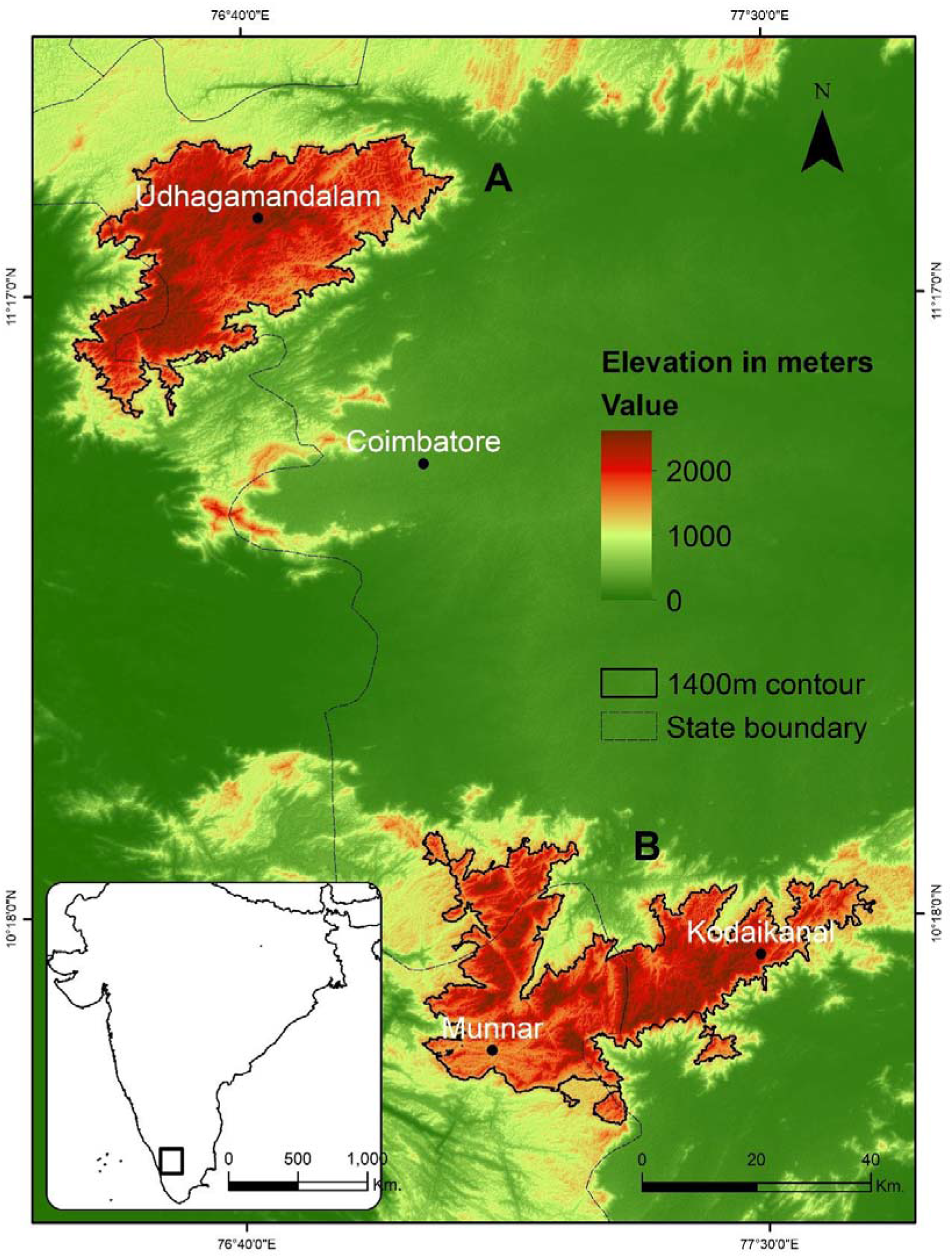
Location map of central shola sky islands in (A) the Nilgiris and, (B) the Palani and Anamalai Hills

### 2.2. Satellite images acquisition and pre-processing

We downloaded cloud-free RapidEye multispectral images from Planet Labs (https://www.planet.com/products/planet-imagery/). Satellite images were obtained in the dry season (January to March 2019) when the water vapour and cloud cover were very low in the atmosphere. The RapidEye satellite images have 5 m spatial resolution and five spectral bands: blue (440 - 510nm), green (520 - 590nm), red (630 - 685nm), red edge (690 - 730nm) and NIR (760 - 850nm).

We converted the RapidEye radiance images into surface reflectance using FLAASH (Fast Line-of-sight Atmospheric Analysis of Hypercubes) module in ENVI v5.2. We used the following FLAASH input parameters for atmospheric corrections - sensor type, pixel size, acquisition date and time, scene centre location, which were all obtained from the metadata file of the satellite images; ground elevation was obtained from the field. The value for sensor altitude was taken from the European Space Agency (https://earth.esa.int/web/eoportal/satellite-missions/r/rapideye). We used a value of 100 km to parameterize initial visibility for each satellite scene because we had acquired images for the dry season only and the weather conditions were clear (Arasumani et al., 2019). The atmospheric model was considered ‘tropical’ because of the presence of high water vapor content in tropical montane habitats (as in Arasumani et al., 2019). The aerosol model was specified as ‘rural’ because the study area was far away from the urban and industrial sectors (as in Arasumani et al., 2019).

### 2.3. Image classification

We used the object-oriented classification technique for image classification using eCognition (Trimble-Geospatial). We performed the following steps for the image classification, (a) segmentation of images using the multi-resolution algorithm, (b) training the object-based classifier using ground truth points, (c) object-based classification using Random Forest (RF), (d) manual editing and (e) accuracy assessment.

#### 2.3.1 Image segmentation

Image segmentation was performed using the multi-resolution algorithm, which reduces the heterogeneity and increases the homogeneity of classified objects. In this method, ‘objects’ are identified using topological and geometric information and segmented accordingly. The classification of uniform land cover features was conducted using an iterative method with parameters of scale, shape, and compactness. The scale parameter is a combined estimate of spectral and shape heterogeneity, and indirectly influences the size of the image objects. Larger scale parameters, for instance, permit a high level of heterogeneity, generating larger objects. The shape is used to define the ratio of spectral homogeneity to spatial shape. Lower shape values assign more weight to spectral values and higher values indicate more importance of shape over spectral values. Compactness is used to identify the objects with compact vs complex shapes. The values for shape and compactness range from 0 to 1. We used a value of 20 for scale for the segmentation, as we were able to discriminate small elements at this value. We used a value of 0.2 for shape and 0.5 for compactness (after iterating values between 0 and 0.8) as these were observed to produce the highest classification accuracy.

Our initial objective was to identify the invasion front and sites with isolated mature exotic trees for grassland restoration. After extensive fieldwork, we realised that invasion had various alternate configurations only some of which were detectable with remote-sensing images alone. Additionally, we found that multiple species of acacias, pines, and eucalyptus were often found together. Since land managers would treat all these species as exotic invasives for restoration efforts, and discriminating species at the sapling stage using remote sensing images is challenging, we combined these three species into one exotic invasive tree class. All categories of potential restoration sites in this study could thus have a combination of these exotic invasive trees. With this in mind, we arrived at the following four categories of invaded habitats using a combination of remote sensing and field data, that could be targeted for restoration:

##### Category 1: Lightly Invaded Grasslands (LIG)

These are large grasslands that have young (small) invasive trees (Fig. 2A). This area most likely constitutes the invasion-front—an area where these invasive exotic trees are moving into the grasslands. We were able to distinguish these lightly invaded grasslands from the mature exotic trees using RapidEye images with the object-oriented classification described above.

**Fig. 2.**
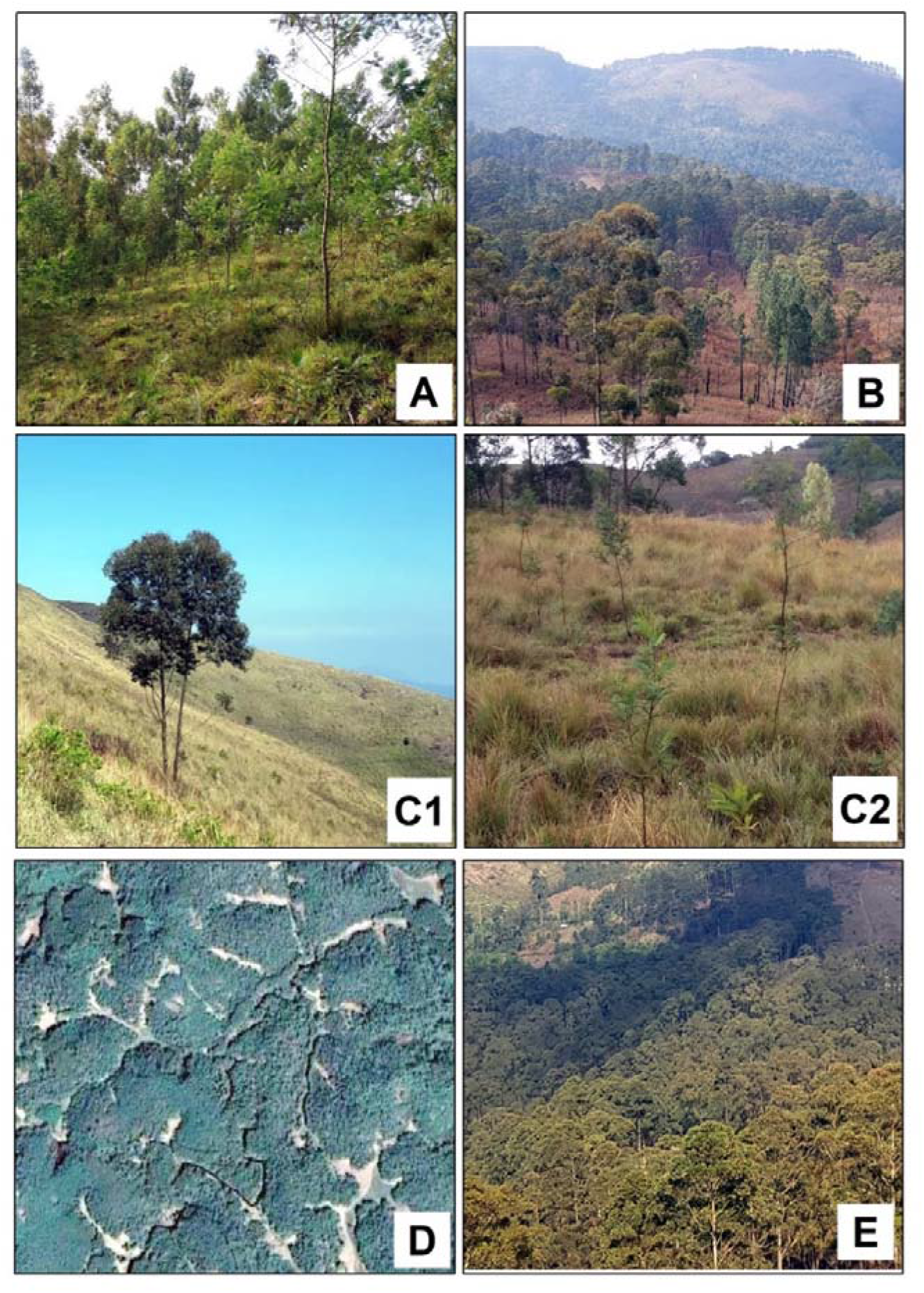
Photographs of potential grassland restoration sites classified as (A) lightly invaded grasslands (LIG), (B) sparse mature exotic tree stands with grass cover (SMG), (C1) isolated exotic trees and (C2) sparse saplings in the grasslands (ITG), (D) fragmented grasslands enveloped by mature exotics trees (GET) and, (E) Mature exotics trees (MET)

##### Category 2: Sparse mature exotic tree stands with grass cover (SMG)

In some areas, grasses persist underneath large, but sparse mature exotic tree stands (Fig. 2B). We detected these patches also using a combination of RapidEye images and object-oriented classification. It may be noted that only a few grass species of the full community of grasses persist under the canopy of mature trees (due to the presence of high shade). Nevertheless, restoration practitioners in the landscape felt that this would be a better starting point for restoration than areas devoid of any natural grasses. Therefore, we retained this landscape type as a separate category.

##### Category 3: Isolated exotic trees and sparse saplings in the grasslands (ITG)

In some areas, we found isolated, but mature exotic trees (Fig. 2C1) and isolated exotic saplings (Fig. 2C2) in the grasslands. The removal of these is critical to prevent them from acting as sources, or in the case of the saplings maturing into future sources, for further invasion. We had difficulty identifying these categories using our automated classification procedures and satellite imagery alone due to the limited spatial and spectral resolution of the imagery. We mapped this category using extensive fieldwork and onscreen digitization from the RapidEye images with a combination of high-resolution Google Earth images. At some locations, native species like Rhododendron may have been identified in this category. The odds of this though are low, since that the shape of a Rhododendron tree is very different from a Eucalypt, pine or acacia.

##### Category 4: Fragmented grasslands enveloped by mature exotics trees - (GET)

Some small grasslands have been enveloped by exotic trees (Fig. 2D). We found these persistent grassland patches in marshes, on hillocks and near streams which had not yet been invaded. Although these were small, we found several grassland-endemic species in these fragmented patches and suggest that these should be included as potential restoration sites. The detection of such patches was the same as for any grasslands. Post-image-classification, we converted such intact, yet fragmented grasslands to the GET category using ArcGIS 10.5. Although these are technically grasslands, management of these patches would have to be different, as restoration between sites can prove to be very fruitful given the persistence of native flora and fauna in this landscape.

In addition to these, we also delineated a fifth category comprising vast areas of mature invasive exotic trees in the landscape i.e. Mature exotic trees (MET; Fig. 2E). We identified MET using RapidEye images and object-oriented classification. We retained MET as a separate class as we anticipate adverse impacts on the soil and other life forms if these are removed rapidly.

We trained the object-based Random Forest (RF) classifier applying ground-truth GPS points of different land covers. The training samples were collected using a field-based method based on systematic sampling for intact grasslands, forest, LIG, SMG, ITG, GET, MET, and water bodies.

The GPS points of training samples were collected using Trimble Juno 5; the GPS accuracy was less than RapidEye image pixel size. We collected 1342 GPS points (Appendix - Fig. 1) from February 2018 to August 2019—approximately the same period as the satellite images—for image classification.

At times, the SMG and MET categories were misclassified into the LIG. We resolved these using manual editing tools in eCognition after extensive field work. We also had a problem with identifying ITG using satellite images alone. This may be due to the spatial and spectral resolution of the imagery used. Since it is necessary to identify these areas and restore them by removing these isolated trees, we used a hybrid approach to identify these. For ITG, all the intact grasslands, which are typically near large stands of exotic trees that we detected from images, were manually checked using Google Earth and RapidEye images, combined with extensive fieldwork. The polygons were then manually edited and corrected using eCognition (Trimble-Geospatial) software.

### 2.4. Accuracy Assessment

The accuracy of the Random Forests (RF) classified map was calculated using ground truth points. We created 350 random points on the classified map and visited these locations to assess the accuracy of the map. We calculated the accuracy using the confusion matrix (Congalton, 1991) in ERDAS IMAGINE (Imagine, 2014).

### 2.5. Identification of grassland restoration sites in the different protect regimes

We used protected areas and range boundaries to estimate the extent and area of each protected regime. We clipped the grassland restoration sites layer with the forest administrative boundaries. The shapefiles containing the boundaries for National Parks, Wildlife Sanctuaries and Reserved Forest were obtained from the Tamil Nadu and Kerala forest departments. The area of grassland restoration sites of each protected region was calculated using ArcGIS v10.5 (ESRI, 2001).

### 2.6. Predicting areas of invasion expansion with logistics regression modeling

We proposed to assess landscape characteristics associated with recently invaded areas. We used a logistic regression modeling approach to evaluate exotic tree invasion into the grassland (i.e. the LIG class). We used the LIG class only since this can be detected using satellite images. The other classes in this study included some manual digitization, which may skew the results and might not be replicable. Our ability to predict exotic tree invasion in the grasslands (i.e. LIG) is based on different independent variables related to topography and the environment and may help identify regions of future spread. We choose the dependent variables (Appendix - Fig. 2) by generating Boolean images indicating the absence of invasives - ‘0’ (including intact grasslands, ITG and GET), and ‘1’ - the presence of lightly invaded exotic plants (LIG).

The independent variables were selected using a literature search and expert consultation. We used the following independent variables: 150 m × 150 m moving window of exotic trees (Appendix - Fig. 3A & B), Topographic Roughness Index (Appendix - Fig. 4A & B), Curvature (Appendix - Fig. 5A & B), fire frequency (Appendix - Fig. 6A & B), and fire intensity (Appendix - Fig. 7A & B). We assumed that the presence of mature trees will impact new invasion and chose a 150 m × 150 m moving window based on our previous study (Arasumani et al., 2018). We included TRI as previous analyses (Arasumani et al., 2018) and current observations showed that steep slopes are resistant to invasion. Similarly, fire frequency and fire intensity have been known to impact invasiveness and we wanted to test this in the Western Ghats landscape (Van Wilgen and Richardson, 1985).

We used focal Statistics with SUM function in ERDAS IMAGINE (Imagine, 2014) to produce 150 m × 150 m moving windows around mature exotic trees. We derived TRI from the ASTER GDEM. We downloaded the MODIS fire intensity and fire frequency data (2014 Dec–2018 Dec), with a spatial resolution of 1000 m and 500 m, from the United States Geological Survey (USGS - https://lpdaac.usgs.gov/product_search/). The dependent and independent variables were resampled to a 30 m spatial resolution and converted into the same projection system. All independent variables were rescaled from 0 to 1.

We used the Akaike information criterion (AIC) to select the best model. We used 80% of random sample data to run a logistic regression model and retained the remaining (20%) to validate the model. We calculated the accuracy of the predicted logistic regression models using AUC/ROC (Area Under Curve/Relative Operating Characteristic). The logistic regression modeling analysis was conducted using R (Appendix 2; Team, 2019) and ArcGIS v 10.5 (ESRI, 2001).

## 3. Results

The overall classification accuracy of intact grasslands, shola forest, LIG, SMG, ITG, GET, MET, and water bodies was 95.43, and the Kappa coefficient was 93.64. Our study identified an area of 254 sq. km. for grassland restoration, 362 sq. km. of intact montane grasslands for conservation, and 606 sq. km. of exotic mature exotic tree stands in the Nilgiris (Table 1, Fig. 3A), Palani Hills and Anamalais (Table 1, Fig. 3B). Of the 254 sq. km., identified areas of restoration, 113 sq. km. were isolated exotic trees and saplings in the grasslands (ITG), 55 sq. km. contained sparse mature trees with grasses (SMG), 45 sq. km. of lightly invaded plants in the grasslands (LIG) and 42 sq. km. of fragmented grasslands patches enveloped by mature trees (GET). Most of the areas suitable for montane grassland restoration were located in the Nilgiris (126 sq. km.), followed by the Palani Hills (73 sq. km.), and the Anamalais (55 sq. km.). The largest areas for grassland restoration were located in Reserved Forests (87 sq. km.), followed by Wildlife Sanctuaries (60 sq. km.) and National Parks (27 sq. km; Table 2). A detailed description of the results which is relevant for conservation managers is included in the Supplementary Materials (Appendix 1-Results).

**Fig. 3.**
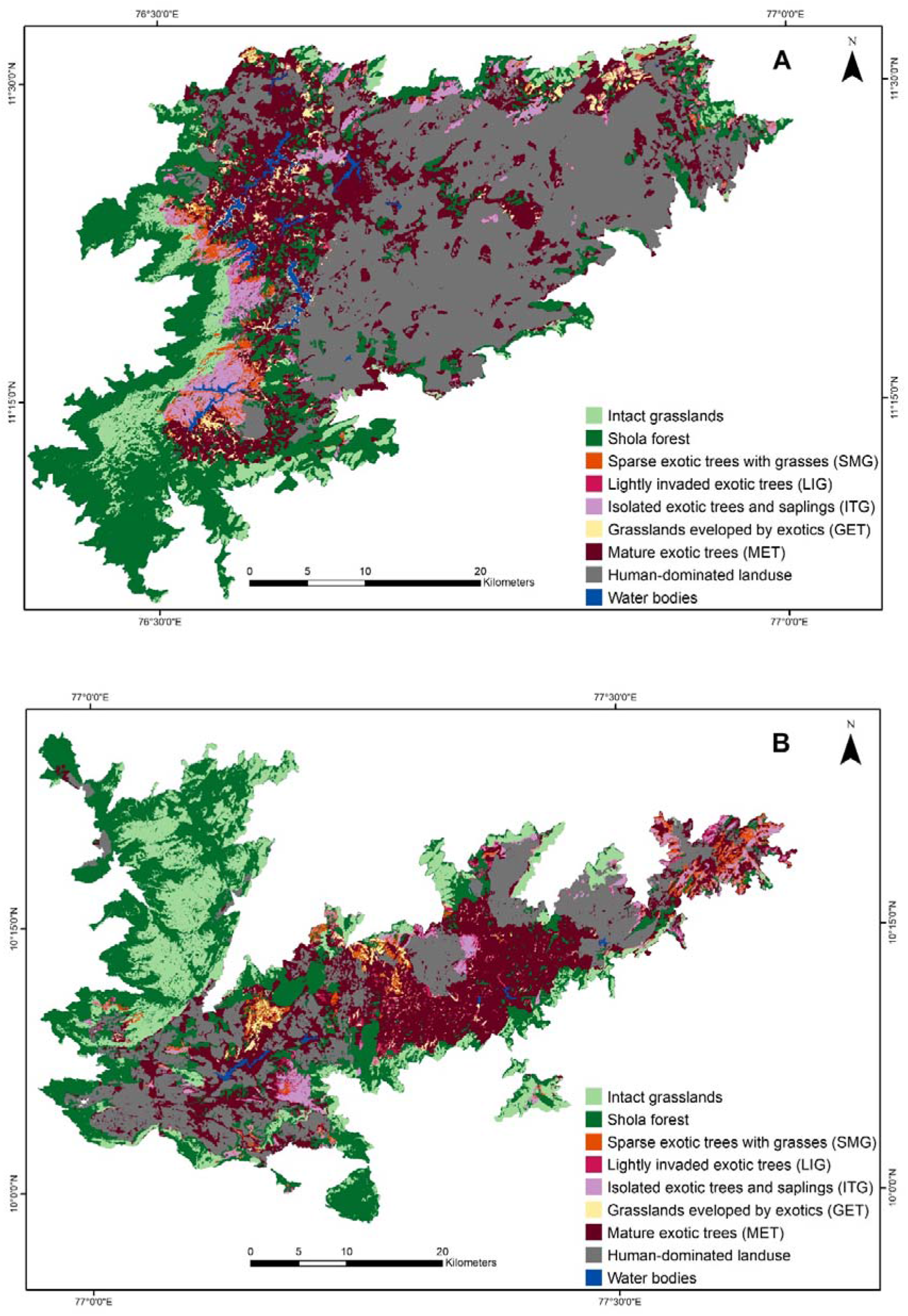
Landuse-landcover map and prioritised sites for grassland restoration in (A) the Nilgiris and, (B) the Palani and Anamalai Hills

**Table 1:** Extent of intact grassland and prioritised sites for grassland restoration in the Western Ghats (sq. km.)

|  | Intact<br>grassland | Priority 1 |  | Priority 2 |  | Priority<br>3 |
| --- | --- | --- | --- | --- | --- | --- |
|  |  | ITG | LIG | SMG | GET | MET |
| Nilgiri Hills | 103.72 | 61.50 | 17.08 | 22.48 | 25.34 | 278.99 |
| Palani Hills | 51.66 | 30.86 | 17.18 | 18.00 | 6.68 | 202.91 |
| Anaimalai Hills | 206.53 | 20.24 | 10.56 | 14.80 | 9.56 | 124.30 |
| Total<br>(central sky<br>islands) | 361.91 | 112.60 | 44.82 | 55.28 | 41.58 | 606.21 |

**Table 2:** Distribution of intact grassland and sites for grassland restoration across protected area management regimes.

|  | Intact<br>grassland | Priority 1 |  | Priority 2 |  | Priority<br>3 |
| --- | --- | --- | --- | --- | --- | --- |
|  |  | ITG | LIG | SMG | GET | MET |
| National Park | 129.37 | 14.18 | 2.56 | 8.69 | 1.85 | 18 |
| Wildlife<br>Sanctuary | 86.38 | 20.59 | 14.93 | 16.35 | 7.97 | 193.32 |
| Reserved Forest | 67.4 | 39.21 | 13.95 | 15.28 | 18.53 | 187.95 |

### 3.1. Factors relating to the exotic invasion front in the montane grasslands

The 150 m × 150 m moving window of mature exotic trees was the best predictor for the invasion front in the grasslands, (β = 3.9, p < 0.001). Areas with low topographic ruggedness value (i.e. smooth terrain) showed higher invasion compared to areas characterised by rugged terrain (β = −1.8, p < 0.001). We did not find any relationship between curvature and invaded areas. We find an inverse relationship between fire frequency and invasion by exotic trees with lower invasion in pixels characterised by a high fire frequency (β = −0.93, p < 0.001). We observed high fire intensity in the invaded areas compared to uninvaded grasslands (β = 1.22, p < 0.001). The AUC_ROC_ of the predicted model was 0.82.

## 4. Discussion

Consistent with our previous efforts (Arasumani et al., 2018; Arasumani et al., 2019) we find that a large part of the montane grasslands has been converted to exotic trees (340 sq. km.). In this study, we identified 254 sq. km. of invaded landscapes for potential restoration to grasslands and an additional 362 sq. km. of intact grasslands for conservation and prevention of invasion in the Western Ghats. In the Western Ghats, there is a strong push for restoration of native habitats, both from forest departments and local communities, driven by issues of water security and biodiversity conservation. A public interest litigation is also directing the state to remove invasives and restore natural habitats. There are modestly sized candidate sites in the National Parks and Wildlife Sanctuaries which have higher protection, and larger candidate sites under areas with lower protection (i.e. Reserved Forests). We have indicated the spatial extent of these sites for each management unit (i.e. Forest Division and subdivision) in the Supplementary materials.

### 4.1. Why should we restore and conserve grasslands?

Grassland habitats have high species richness and contribute significantly to ecosystem services, but this species richness can be reduced by tree plantations (Koch et al., 2016). Such modified landscapes with invasive alien tree species can have a significant impact on global ecosystem services including water security. In South Africa, invasive exotic species like acacia, pine, and Eucalyptus are thought to have considerably reduced soil water and streamflow. Likewise, Dehlin et al. (2008) observed that invasive pines were impacting nutrient levels and soil water in New Zealand. Many exotic trees act as ecosystem transformers and induce changes in the structure and function of the ecosystems they have invaded (Richardson and Higgins, 2000).

Our study indicates that more than half the original extent of grasslands have been lost threatening the taxa that these habitats support. There is also evidence that anthropogenic climate change linked increases in temperature in the future are likely to support exotic tree expansion into the montane grasslands. Further evidence from field-based experiments suggests that the rates of expansion of the exotic, invasive *Acacia spp.* is likely to be much higher than that of native trees (Joshi et al., 2020). There is, thus, an urgent need to control grassland invasion and undertake conservation and restoration efforts.

### 4.2. Prioritizing between different patterns of invasion, and restoration sites

Nearly 10 years ago, Le Maitre et al. (2011) noted that millions of dollars had been spent on the control and removal of IAS including acacias. Given the expense in controlling invasive trees such as acacias that have large and long-lived seed banks (Le Maitre et al., 2011), restoration, we join numerous researchers (e.g. Souza-Alonso et al., 2017) in recommending the prioritisation of restoration efforts.

Our study finds that there are differences in landscape patterns of invasion by exotic trees. Based on this, we categorized them into four distinct types of invaded habitats. The restoration strategies in each of these categories, and consequently the ecological complications may be different in each of these types. Further, the challenges in detecting (and eventually monitoring) these different invaded habitats using remotely sensed data are also unique. We propose that the categories of potential restoration sites described in this study be used as a starting point to prioritize restoration activities.

#### Priority 1

We recommend that the first restoration efforts be focussed on the 113 sq. km. of grasslands with isolated exotic trees (ITG) and the 45 sq. km. of lightly invaded grasslands (LIG). These isolated trees in grasslands (ITG), or young plants in grasslands (LIG) we believe can be removed with relatively low effort, and the area of grasslands recovered, although not the largest in the landscape, is still quite high. No additional planting of grasses would be required in these areas, reducing the costs of the restoration effort. The only intervention required to get the system back to a healthy state may be the removal of these invasive elements in small quantities. These restoration efforts can also help contain the rapid spread of invasives to uninvaded areas, with a potentially high return, for a relatively low investment (of man-hours and money). Some of our LIG sites contained native grassland trees (e.g. *Rhododendron arboreum nilagiricum (Zenker)*), but the expertise of restoration workers knowledgeable with the landscape should be sufficient in retaining these trees during restoration activities.

#### Priority 2

Sparse mature trees that still have grass on the floor (SMG) occupy approximately 55 sq. km. which, with fragmented grasslands patches (smaller than 1 ha) enveloped by mature trees (GET; spread over 42 sq. km.) could be the second priority for restoration. These areas will require higher effort and investment since mature trees have to be removed. This will also necessitate consideration of other ecological impacts of exotic tree removal. Further although grasslands still exist in these areas, additional planting of grasses can substantially improve the outcome of restoration efforts. We believe that the restoration efforts in GET sites will connect these to large grasslands so that native flora and fauna can continue to persist in these patches. Nevertheless, we recognize that it will be challenging to target all GET fragments and recommend that the choice of the fragments be determined by logistics, including the proximity to the nearest large grassland, and the maximum grassland area that will be connected by each effort.

Finally, there are 606 sq. km. of mature exotic trees in this landscape (MET), an area more extensive than any other category, higher even than the intact grasslands remaining in the landscape. While we do not rule these areas out as restoration sites, we highlight below the significant challenges likely to be encountered in tackling invasion at these sites.

Globally, forest administrators and managers are facing a challenge in restoring grasslands by removing mature exotic trees; this is especially true for *Acacia mearnsii* (Cheney et al., 2019; Le Maitre et al., 2011). Researchers in tropical and sub-tropical landscapes have even noted the extensive presence of native trees and their associated native forest-dwelling taxa in these areas (Geldenhuys, 1997; Srimathi et al., 2012). Not all of the 606 sq. km. of mature exotic areas have forest regeneration, and landscape features associated with such regeneration have not been examined. Neither are there any maps on the extent of such regeneration. In areas with such patterns, however, it is clear that native biodiversity, including several threatened, endemic taxa exist (personal observations of authors). Removing mature exotic trees will negatively affect these species, and each location must be considered separately for potential ecological impacts.

In some areas, large-scale removal may also create other ecological issues. Much of the MET areas do not have a grass layer on the floor. Not only does the removal of mature trees render these locations prone to soil erosion, but grass species will have to be planted on a large-scale to restore the grassland. This, in turn, requires the establishment of capacity at multiple levels. First, a knowledge capacity on native grasses needs to be established to guide the choice of grass species to be restored. Then, nurseries have to be established so these species can be propagated and supplied to various restoration sites. Importantly, substantial and sustained effort will be needed for many years to prevent exotic trees from re-establishing through the existing seed bank.

These efforts are best attempted at smaller areas and on an experimental basis, as has already been initiated in the states of Kerala (Pattiyangal; 20 ha, Pazhathottam; 10 ha) and Tamil Nadu (Kodaikanal and Poomparai; 25 ha each). A thorough documentation of the processes, costs and challenges in restoration at these sites, will aid in developing restoration plans at a landscape scale.

### 4.3. Where are we likely to see more invasion?

Our analyses of invasion front patterns (LIG areas) suggest that invasion is strongly influenced by the proximity to mature exotic trees (MET). Prioritising action at LIG sites is, therefore, as critical to stop further invasion, as it is to restore grasslands. Our results also indicate that invasion was higher in areas of low topographic roughness (smooth terrain). This may reflect habitat preferences (e.g. soil depth and drainage) for species such as *Acacia mearnsii*, although a precise inference is beyond the scope of our study. Significantly, our results also imply that fire frequency can potentially prevent invasion by exotic trees, a conclusion meriting serious thought by state forest departments that tend to exclude fires from these grasslands. On the other hand, fires were more intense in invaded areas, potentially due to the higher fuel load from the exotic trees. A similar relationship (between fire intensity and grassland invasion) has also been reported in South Africa (Van Wilgen and Richardson, 1985), but more experimental data and observation is required to provide further clarity on these patterns.

### 4.4. The ability of remotely sensed images to detect and monitor landscapes for restoration

This research shows the potential uses of high-resolution satellite imagery (e.g. RapidEye) for invasive species management and identifying areas for grassland restoration. A combination of such high-resolution imagery was very useful in detecting young invasion, but we highlight the critical role of extensive field data that was crucial to inform the identification of additional types of restoration sites. At the time our research was conducted, these were the highest resolution images that were available without incurring significant costs. This is a major consideration for tropical, developing countries where such challenges may lie. We were able to quantify the lightly invaded grasslands and mature trees with grasses, but were unable to identify isolated trees and saplings in the grasslands using automated classification algorithms because of the limited spatial and spectral resolution of the images we used. Perhaps using advanced remote sensing technologies such as LiDAR and hyperspectral data, or hyperspatial data, will improve our ability to detect invasion patterns and prioritise sites for restoration. Nevertheless data availability remains a challenge for developing countries like India, and its applicability needs to be evaluated. Despite these challenges, we believe that remotely sensed data is useful not just for detecting and prioritising restoration sites at landscape scales as described here, but also for monitoring restoration activities and to establish the success of these efforts in the future.

## 5. Conclusion

Both biological invasion and ecological restoration are highly complex phenomena, and there are doubtlessly other parameters that would need to be considered while selecting restoration sites that have not been addressed in our study. For example, Le Maitre et al. (2011) also suggested accounting for acacia seeds banks, fire history (which also emerged as an important predictor of grassland invasion in our study), soil characteristics, and leaf litter before restoration efforts. Our study only examines broad characteristics to provide actionable maps for restoration efforts.

Nevertheless, this study demonstrates the potential for remotely sensed data, supplemented with field expertise, to be used to map potential restoration sites over large (landscape-scale) extents. Our study also brings conservation and restoration focus to grassland habitats that have been historically ignored when compared to the concomitant forests from these landscapes (Arasumani et al., 2019; Joshi et al., 2018). While many studies have attempted to map potential forest restoration sites using remotely sensed data (Liu et al., 2019), this approach is uncommon for mapping sites for restoring grasslands. We hope that our approach to identifying grassland restoration sites can aid similar efforts in other landscapes.

## Acknowledgements

This study was supported by the National Geographic Society (NGS-53244C-18) to V.V. Robin and an IISER-Tirupati postdoctoral research fellowship to Arasumani M. We thank Planet’s Education and Research Program for providing access to RapidEye images. We thank Kerala and Tamil Nadu forest departments for research permits and logistical support. We thank C.K. Vishnudas, Vasanth Bosco, Arundhati Das, Madhura Niphadkar, Bishwarup Paul and Arun Govindarajulu for discussions and feedback. We thank Dhanish R., Meera M.R., Dini Das for GIS and fieldwork; Kartik Shanker and Ravi Bhalla for help with logistics; Eco-Evo lab members at IISER-Tirupati for comments and feedback on the manuscript.

## Supplementary Material

### Appendix

**Appendix - Table 1.** Area of potential sites for grassland restoration and intact grasslands in each of the protected regimes.

| Division | Protected<br>manageme<br>nt regimes | SMG | LIG | ITG | GET | MET | Intact<br>grasslands |
| --- | --- | --- | --- | --- | --- | --- | --- |
| Anamudi Shola NP | NP | 1.42 | 1.07 | 0.00 | 1.27 | 10.92 | 0.93 |
| Eravikulam NP | NP | 0.58 | 0.43 | 0.79 | 0.30 | 3.17 | 73.91 |
| Mukurthi NP | NP | 6.68 | 0.32 | 13.2<br>0 | 0.18 | 2.52 | 34.36 |
| Pampadum NP | NP | 0.00 | 0.74 | 0.19 | 0.10 | 1.39 | 0.00 |
| Grass Hill NP | NP | 0.00 | 0.00 | 0.00 | 0.00 | 0.00 | 17.36 |
| Mathiketan Shola | NP | 0.00 | 0.00 | 0.00 | 0.00 | 0.00 | 1.33 |
| Silent Valley NP | NP | 0.00 | 0.00 | 0.00 | 0.00 | 0.00 | 1.49 |
| Chinnar Wildlife<br>Sanctuary | WLS | 0.00 | 0.07 | 0.08 | 0.00 | 0.00 | 8.21 |
| Kodaikanal wildlife<br>sanctuary | WLS | 13.6<br>1 | 13.2<br>4 | 20.5<br>1 | 5.20 | 171.8<br>4 | 34.57 |
| Kurinjimala | WLS | 2.73 | 1.62 | 0.00 | 2.77 | 20.27 | 1.33 |
| Indira Gandhi<br>(Annamalai) | WLS | 0.00 | 0.00 | 0.00 | 0.00 | 1.19 | 42.28 |
| Bagalkot | RF | 0.48 | 0.41 | 0.61 | 1.30 | 5.31 | 1.89 |
| Chinnar | RF | 0.37 | 0.25 | 0.68 | 0.07 | 0.33 | 11.20 |
| Coimbatore | RF | 0.00 | 0.00 | 0.00 | 0.00 | 0.44 | 0.73 |
| Gudalur | RF | 2.68 | 0.34 | 2.57 | 1.25 | 9.65 | 3.62 |
| Kottayam | RF | 0.00 | 0.06 | 0.11 | 0.00 | 0.00 | 0.20 |
| Malayattur | RF | 0.75 | 0.56 | 0.88 | 0.13 | 0.59 | 10.05 |
| Munnar | RF | 0.37 | 0.12 | 0.00 | 0.29 | 0.90 | 2.12 |
| Nilgiris Division | RF | 9.56 | 11.5<br>2 | 31.6<br>6 | 15.5<br>0 | 164.6<br>3 | 27.75 |
| Theni | RF | 1.05 | 0.70 | 2.71 | 0.00 | 6.11 | 9.84 |

**Appendix - Table 2.** Landuse/landcover statistics (in sq.km)

| **Landcover** | **Nilgiris** | **Palani Hills** | **Anaimalai Hills** | **Total (Central sky islands)** |
| --- | --- | --- | --- | --- |
| Intact grasslands | 103.72 | 51.66 | 206.53 | 361.91 |
| Shola forest | 366.32 | 62.08 | 374.08 | 802.47 |
| SMG | 22.48 | 18.00 | 14.80 | 55.28 |
| LIG | 17.08 | 17.18 | 10.56 | 44.82 |
| ITG | 61.50 | 30.86 | 20.24 | 112.60 |
| GET | 25.34 | 6.68 | 9.56 | 41.58 |
| MET | 278.99 | 202.91 | 124.30 | 606.21 |
| Human-dominated landuse | 569.08 | 130.30 | 178.46 | 877.84 |
| Water bodies | 17.20 | 1.32 | 3.08 | 21.60 |

**Appendix - Fig. 1.**
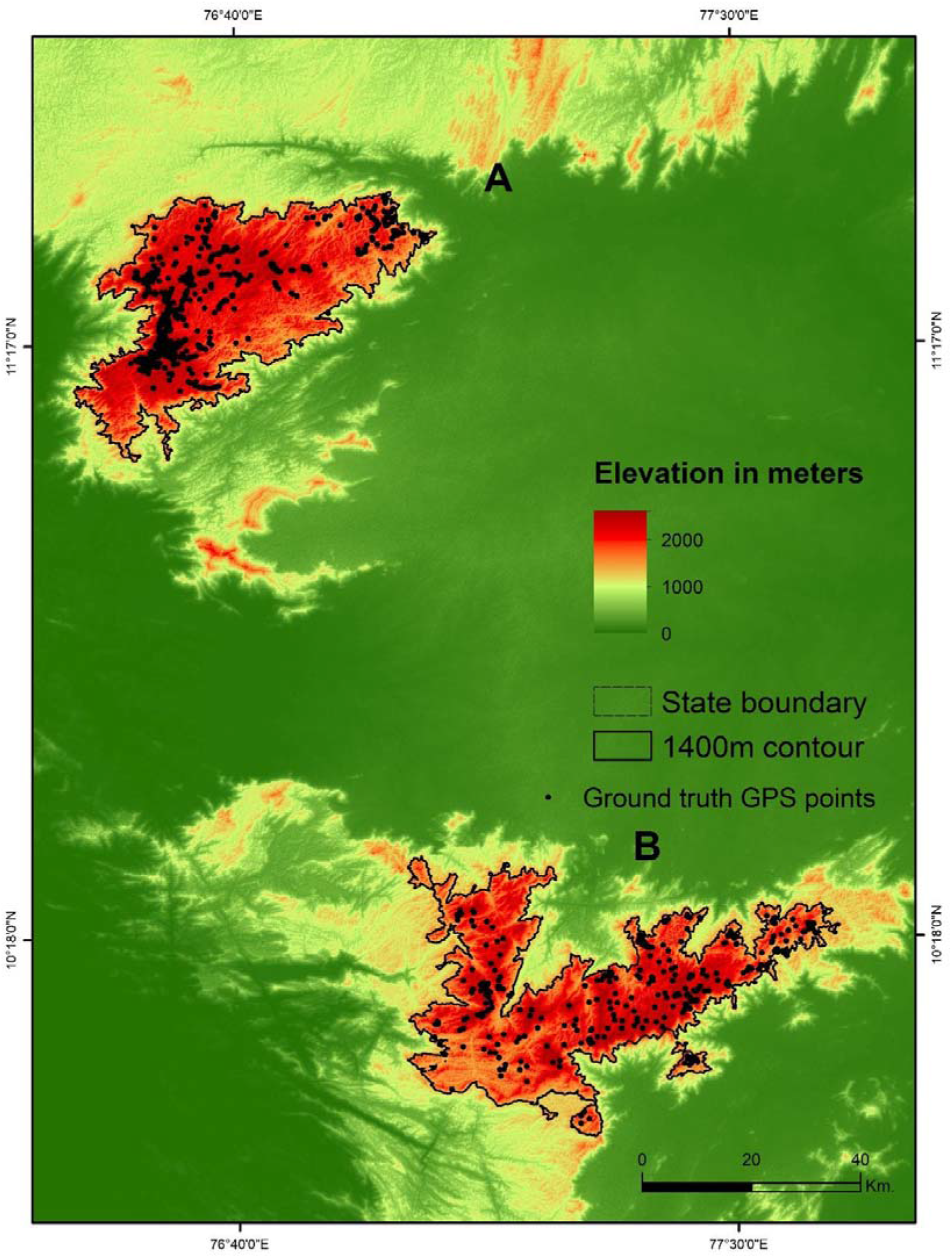
Ground truth GPS points (A) Nilgiris (B) Palani and Anamalai Hills

**Appendix - Fig. 2.**
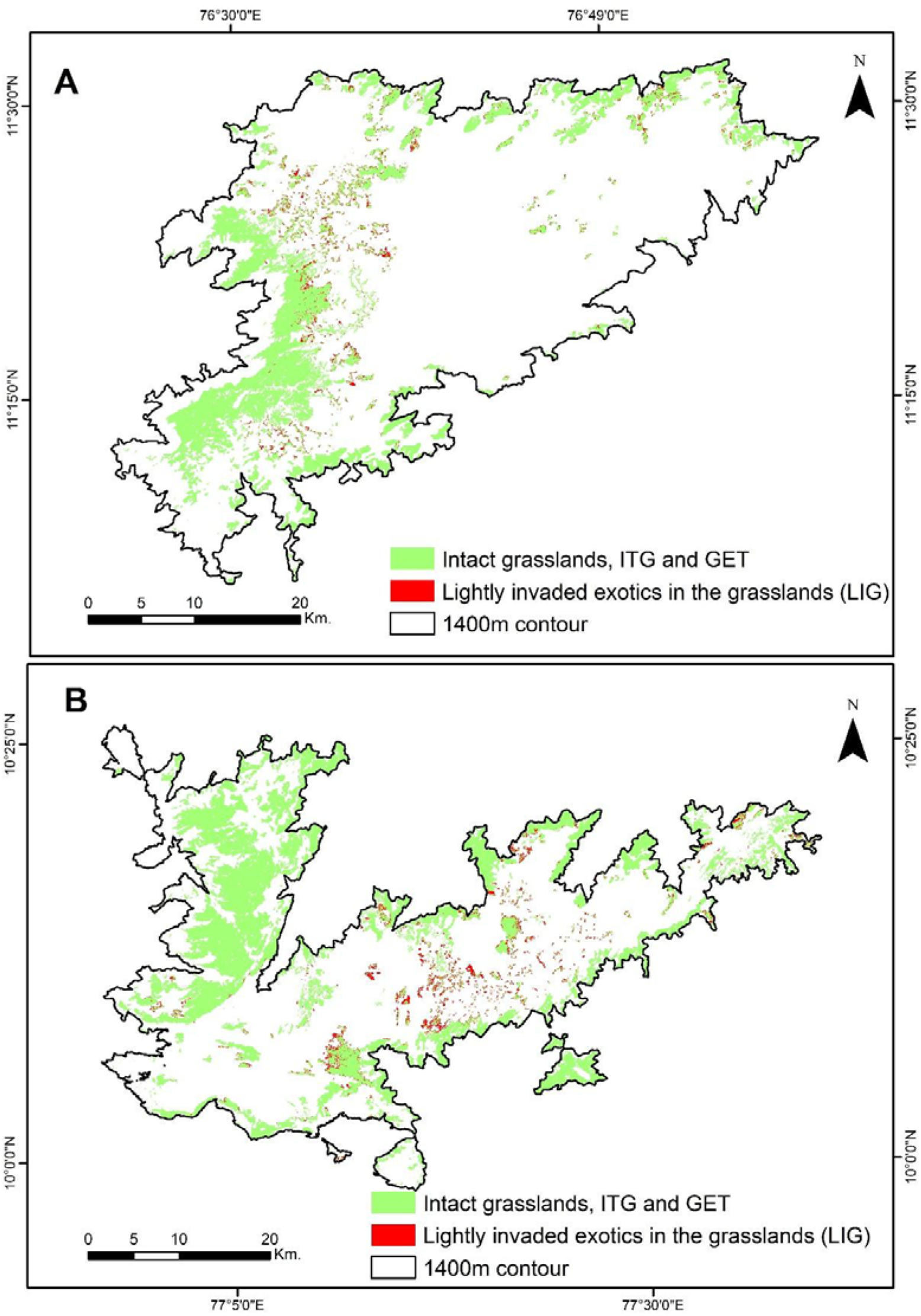
Dependent variable used for Predicting areas of invasion expansion with LRM – (A) Nilgiris (B) Palani and Anamalai Hills

**Appendix - Fig. 3.**
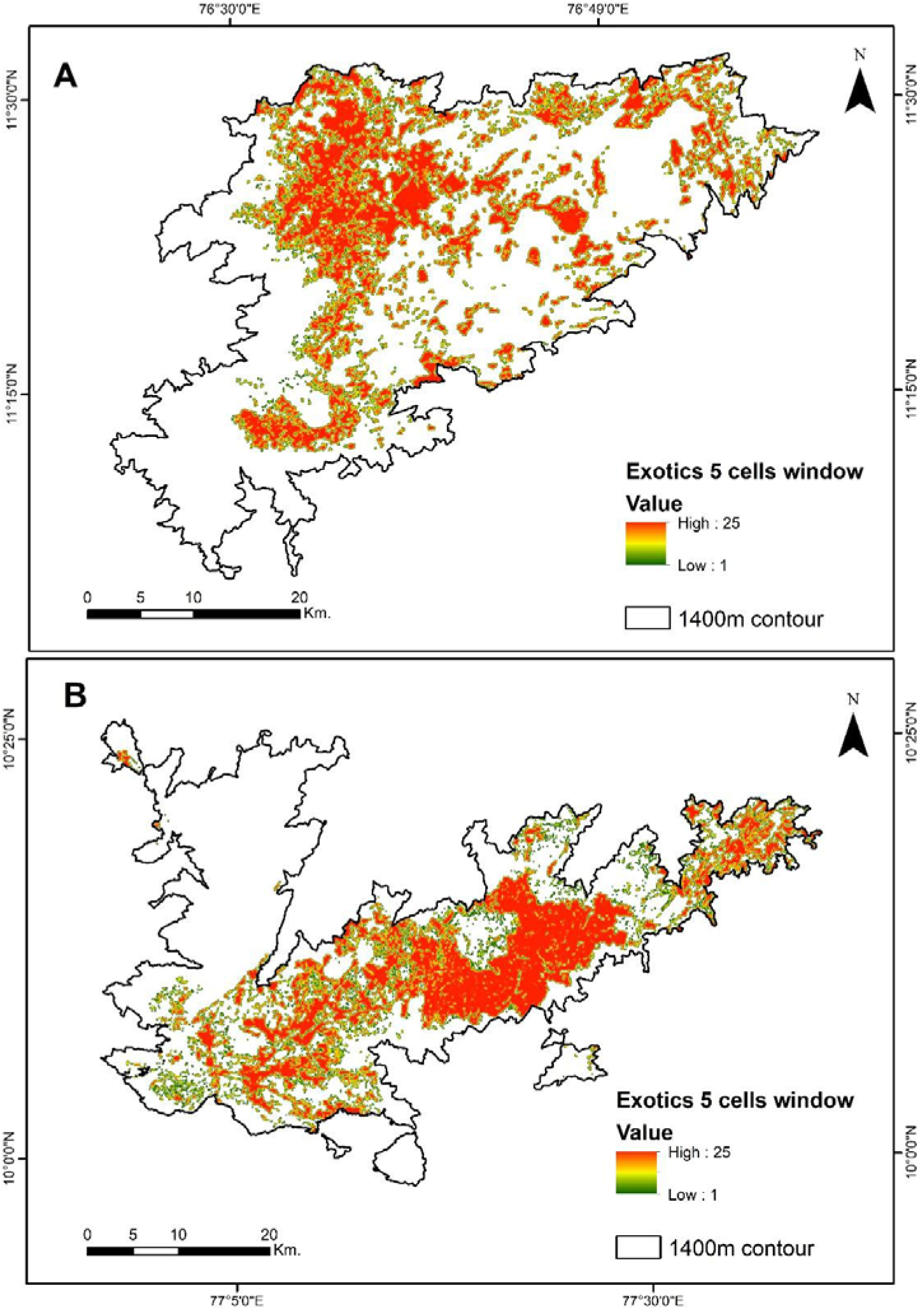
Independent variable used for Predicting areas of invasion expansion with LRM - 150 m × 150 m moving window of exotic trees (A) Nilgiris (B) Palani and Anamalai Hills.

**Appendix - Fig. 4.**
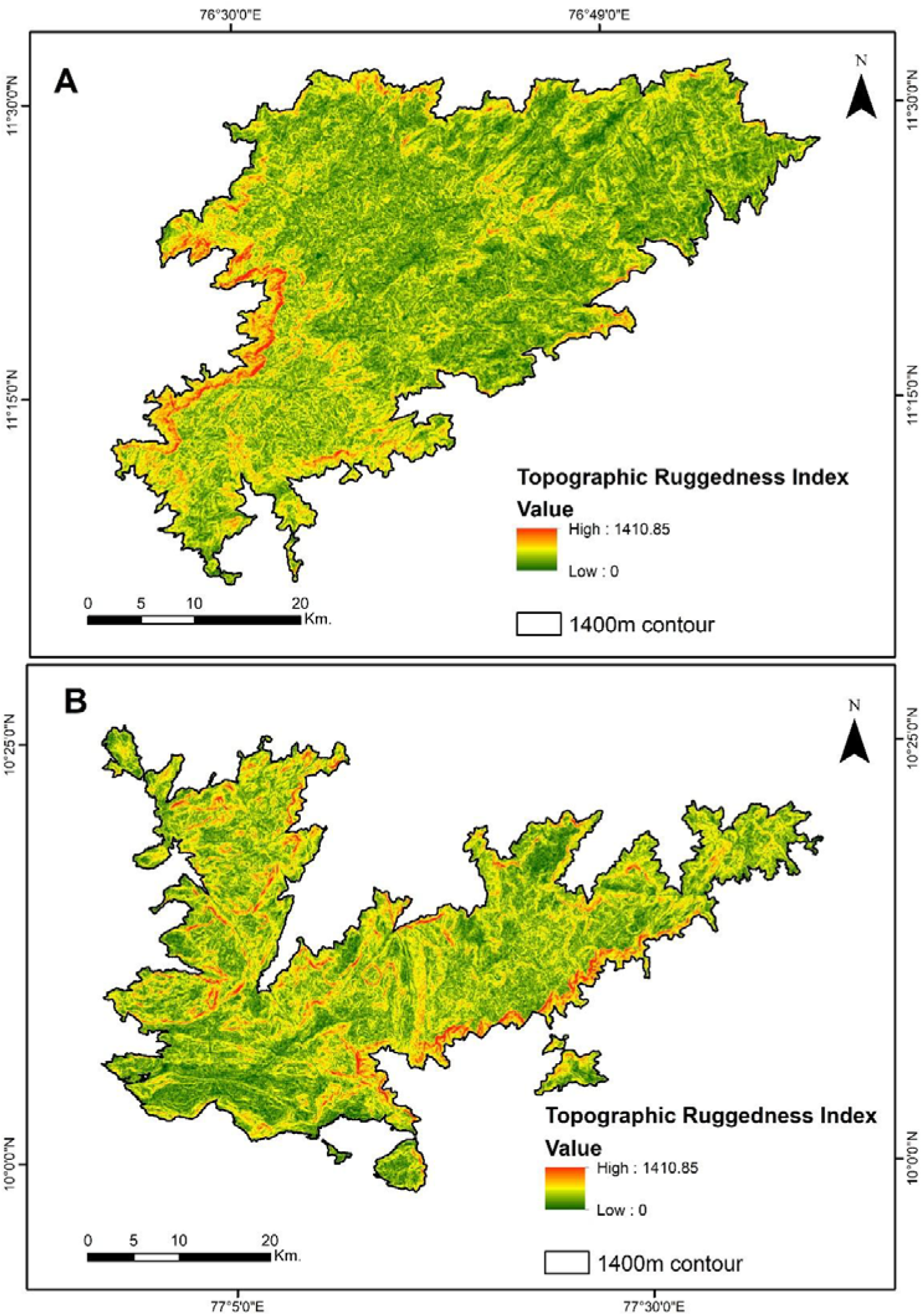
Independent variable used for Predicting areas of invasion expansion with LRM - Topographic Roughness Index (A) Nilgiris (B) Palani and Anamalai Hills.

**Appendix - Fig. 5.**
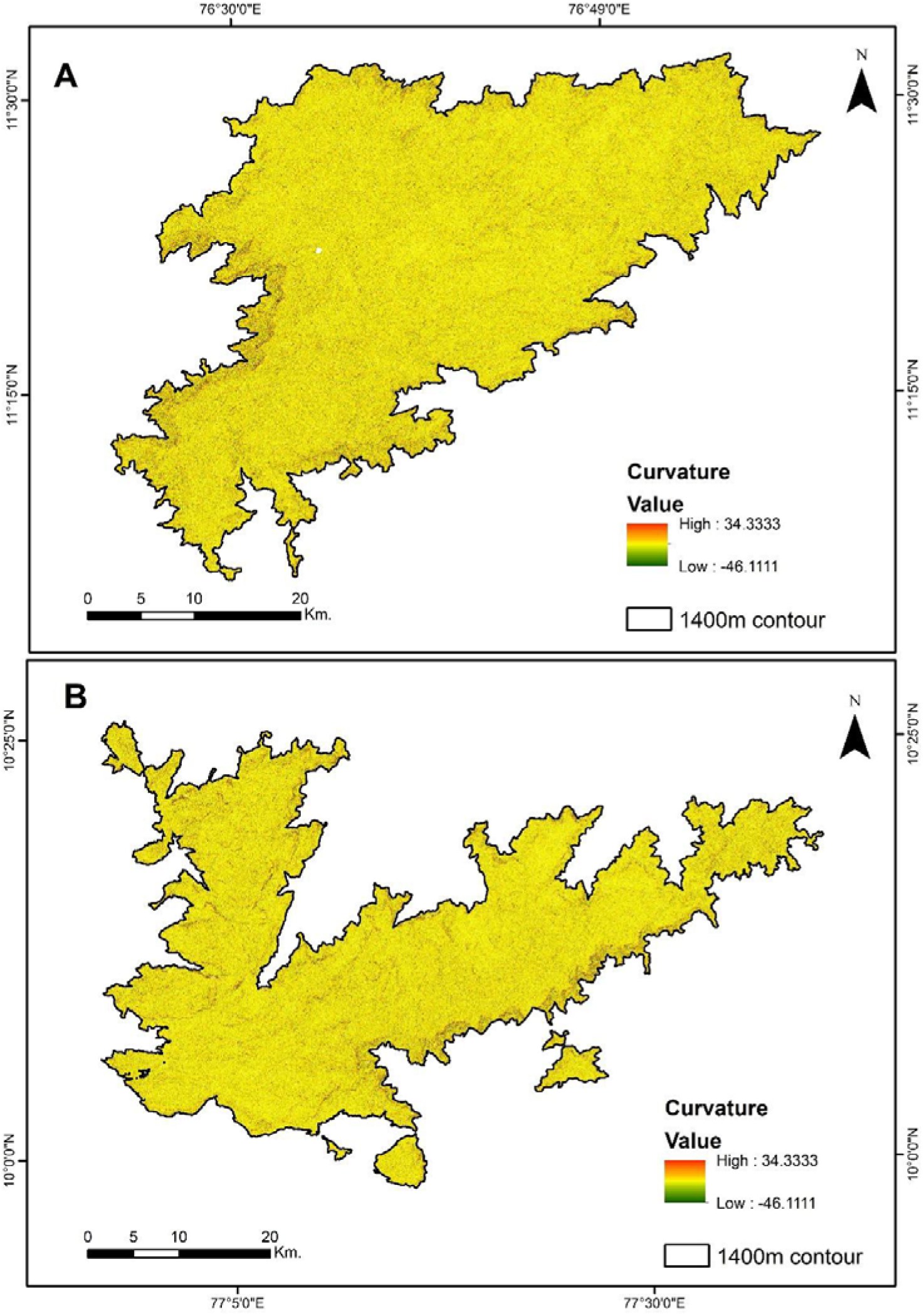
Independent variable used for Predicting areas of invasion expansion with LRM - Curvature (A) Nilgiris (B) Palani and Anamalai Hills.

**Appendix - Fig. 6.**
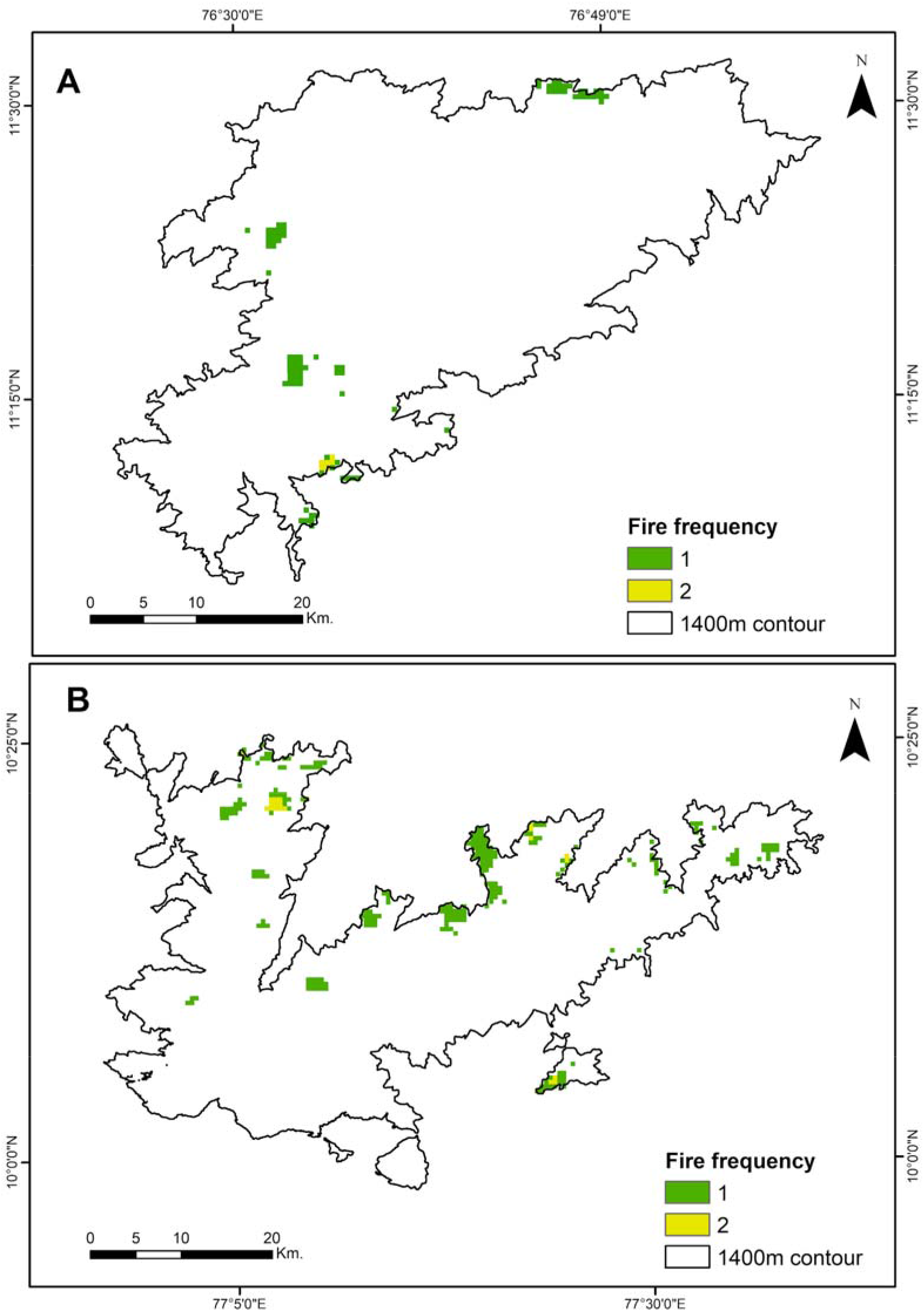
Independent variable used for Predicting areas of invasion expansion with LRM - Fire Frequency (A) Nilgiris (B) Palani and Anamalai Hills.

**Appendix - Fig. 7.**
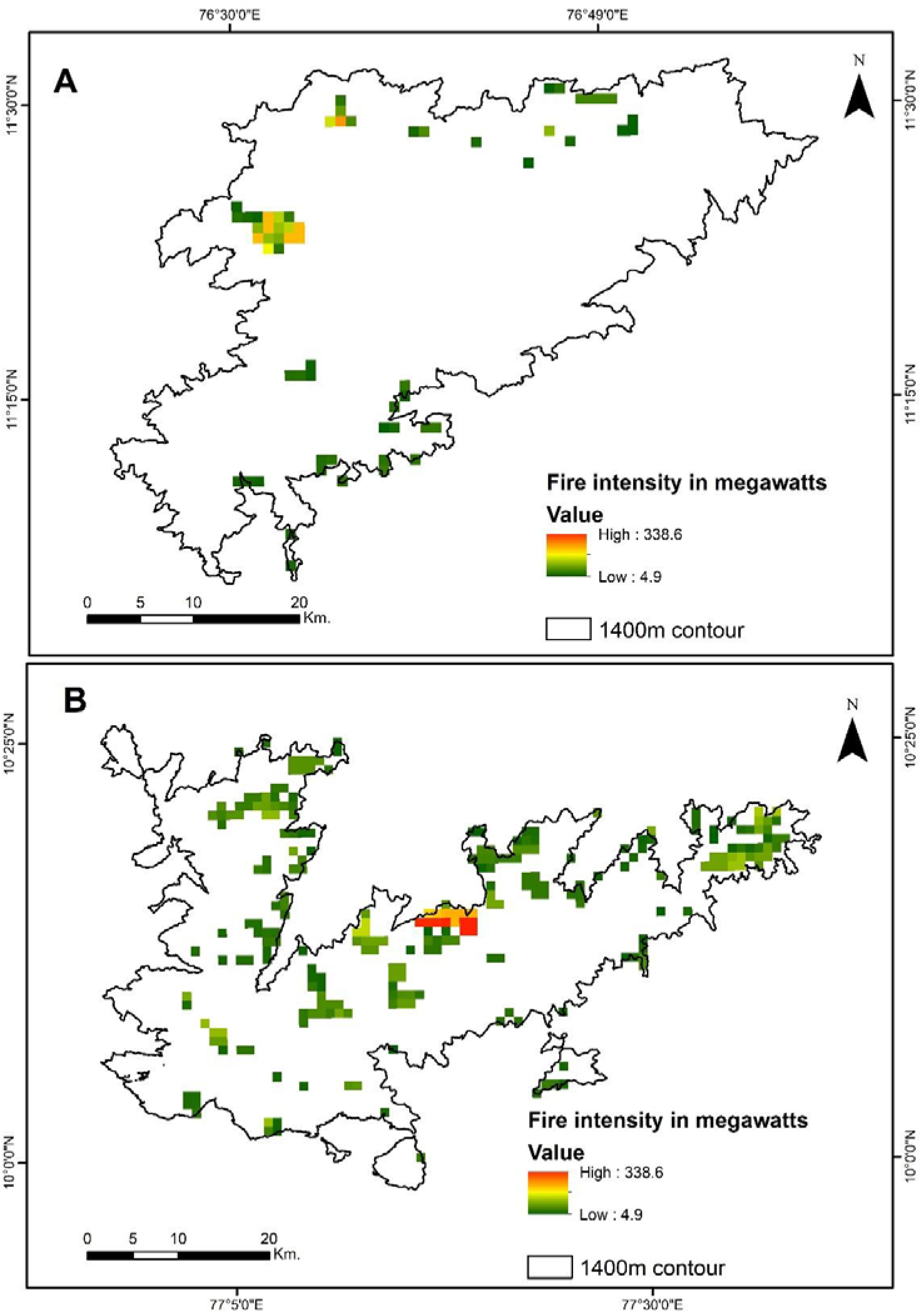
Independent variable used for Predicting areas of invasion expansion with LRM - Fire Intensity (A) Nilgiris (B) Palani and Anamalai Hills.

#### Appendix 1 - Results

Administratively most of the restoration sites were located in the state of Tamil Nadu (199 sq. km.) followed by Kerala (55 sq. km.). We observed that most of the grassland restoration sites were located in the Nilgiri North division (68 sq. km.), followed by Kodaikanal Wildlife Sanctuary (53 sq. km.), Mukurthi National Park (23 sq. km.), Kurinjimala Wildlife Sanctuary (7 sq. km.), Gudalur Forest Division (7 sq. km.), Theni Forest Division (4 sq. km.) and Anaimudi Shola National Park (4 sq. km.).

The largest areas of mature exotic trees are located in the Kodaikanal Wildlife Sanctuary (172 sq. km.) followed by Nilgiri North Forest Division (165 sq. km.), Kurinjimala Wildlife Sanctuary (20 sq. km.), Anaimudi Shola National Park (11 sq. km.), Gudalur Forest Division (10 sq. km.), Theni Forest Division (6 sq. km.), Bagalkot Forest Division (5 sq. km.), and Eravikulam National Park (3 sq. km.).

The largest area of intact montane grasslands were found in Eravikulam National Park (74 sq. km.) followed by Indira Gandhi Wildlife Sanctuary (42 sq. km.), Mukurthi National Park (34 sq. km.), Kodaikanal Wildlife Sanctuary (34 sq. km.), Nilgiris North Forest Division (28 sq. km.), Grass Hills National Park (17 sq. km.), Chinnar Forest Division (11 sq. km.), Malayattur Forest Division (10 sq. km.), Theni Forest Division (10 sq. km.), Chinnar Wildlife Sanctuary (8 sq. km.), Gudalur Forest Division (4 sq. km.), and Munnar Forest Division (2 sq. km.). Very few intact grassland patches were found in other protected regimes.

#### Appendix 2 - R code used for predicting areas of invasion expansion into the grasslands with LRM

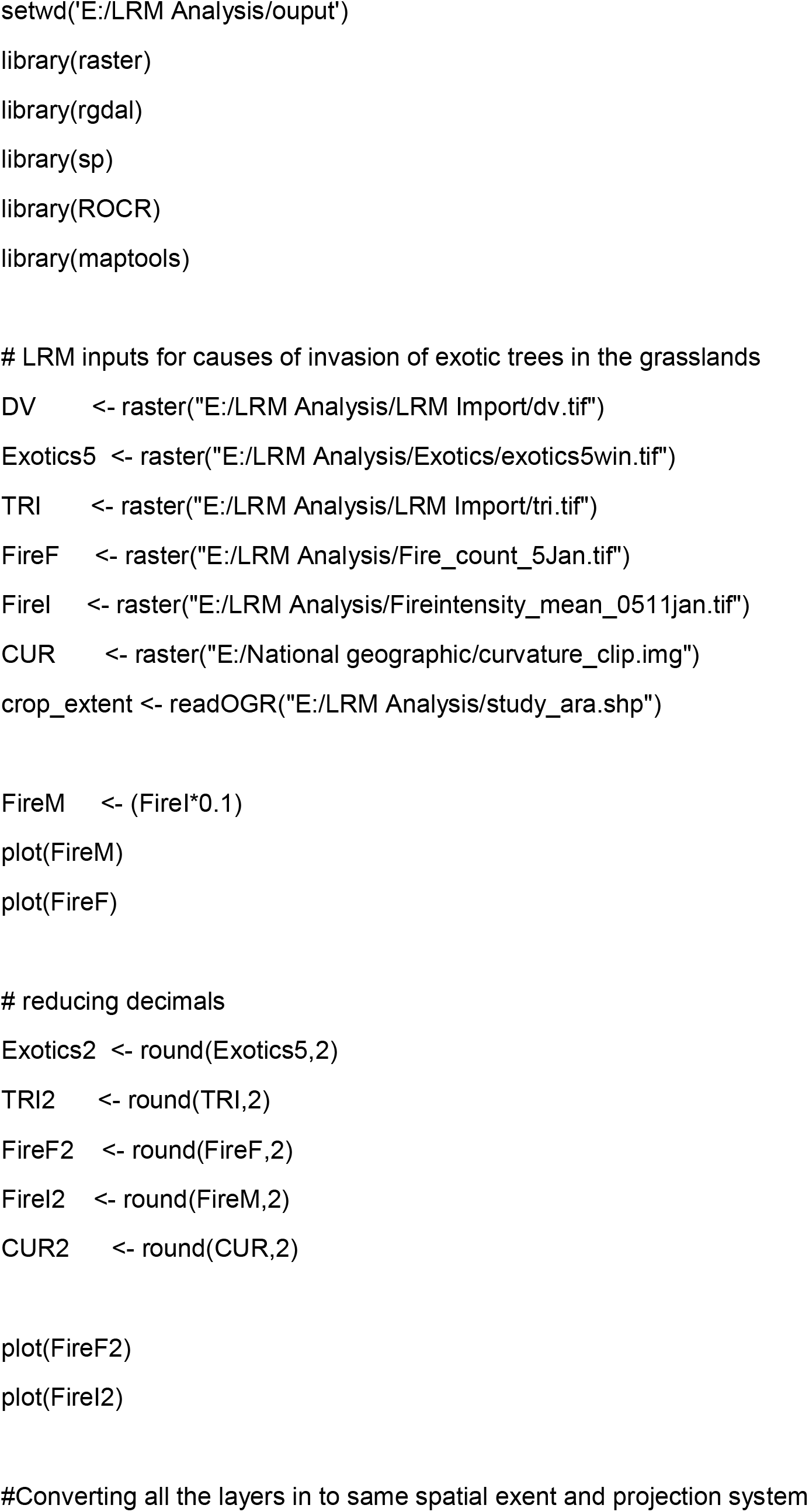
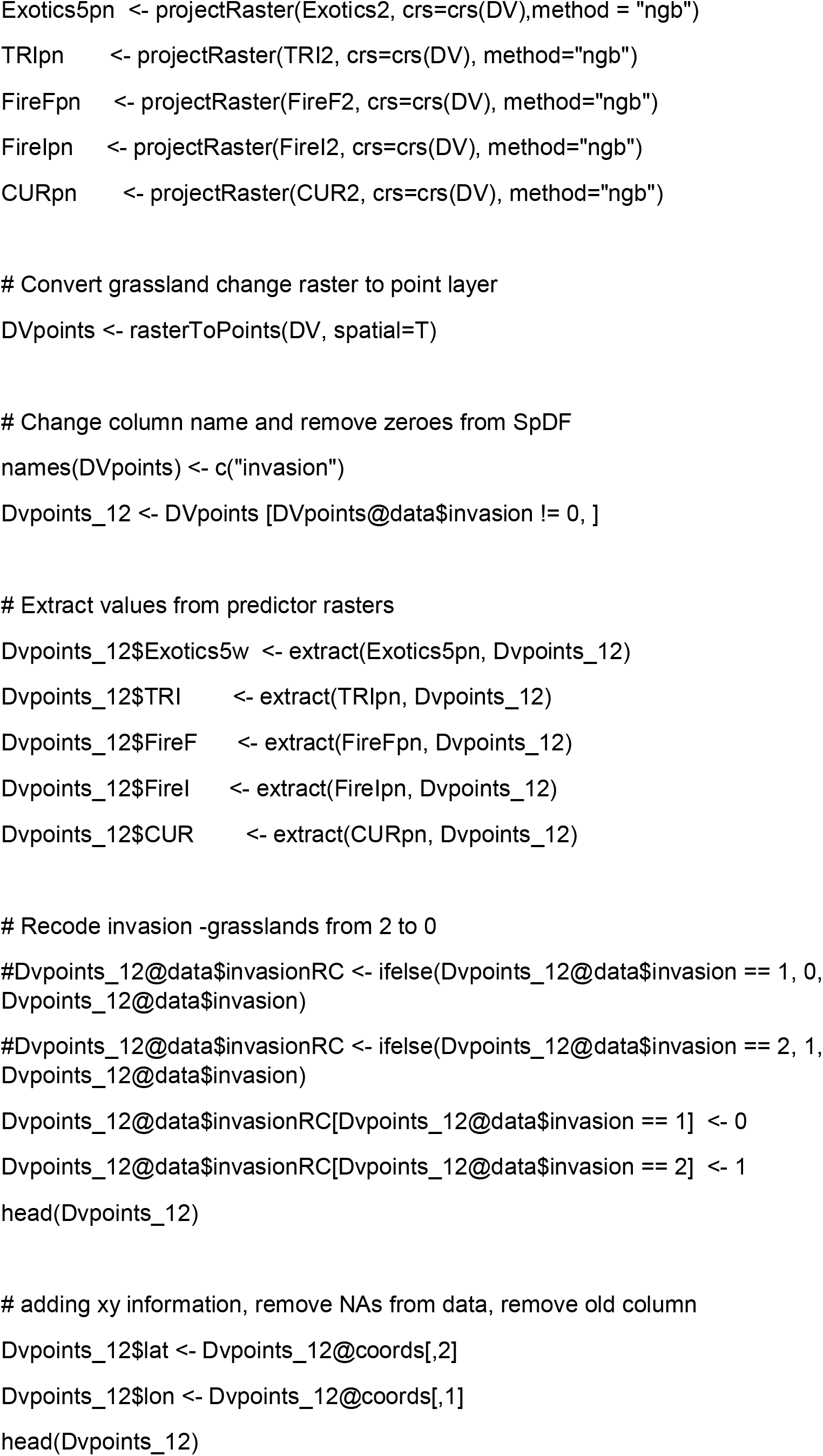
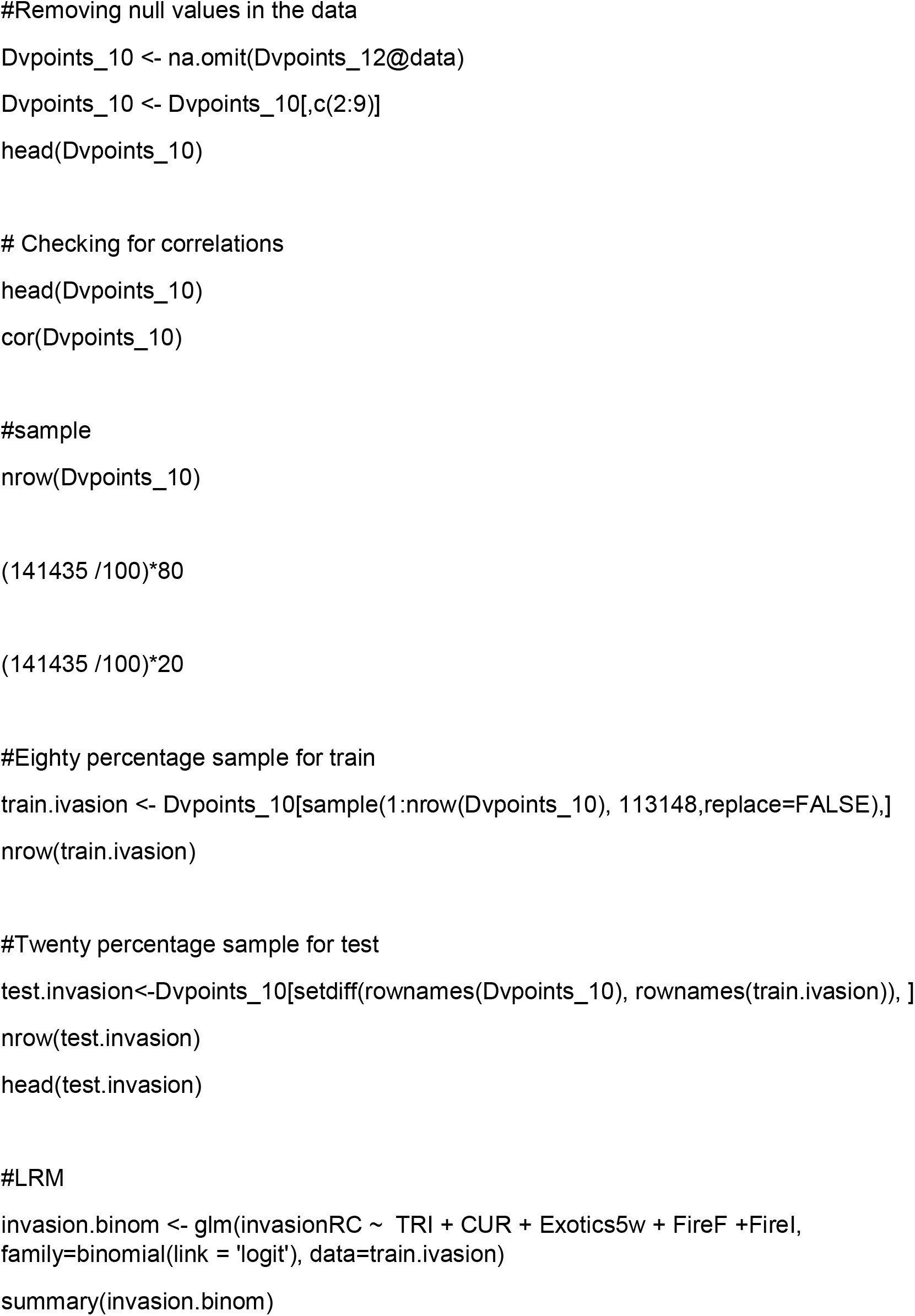
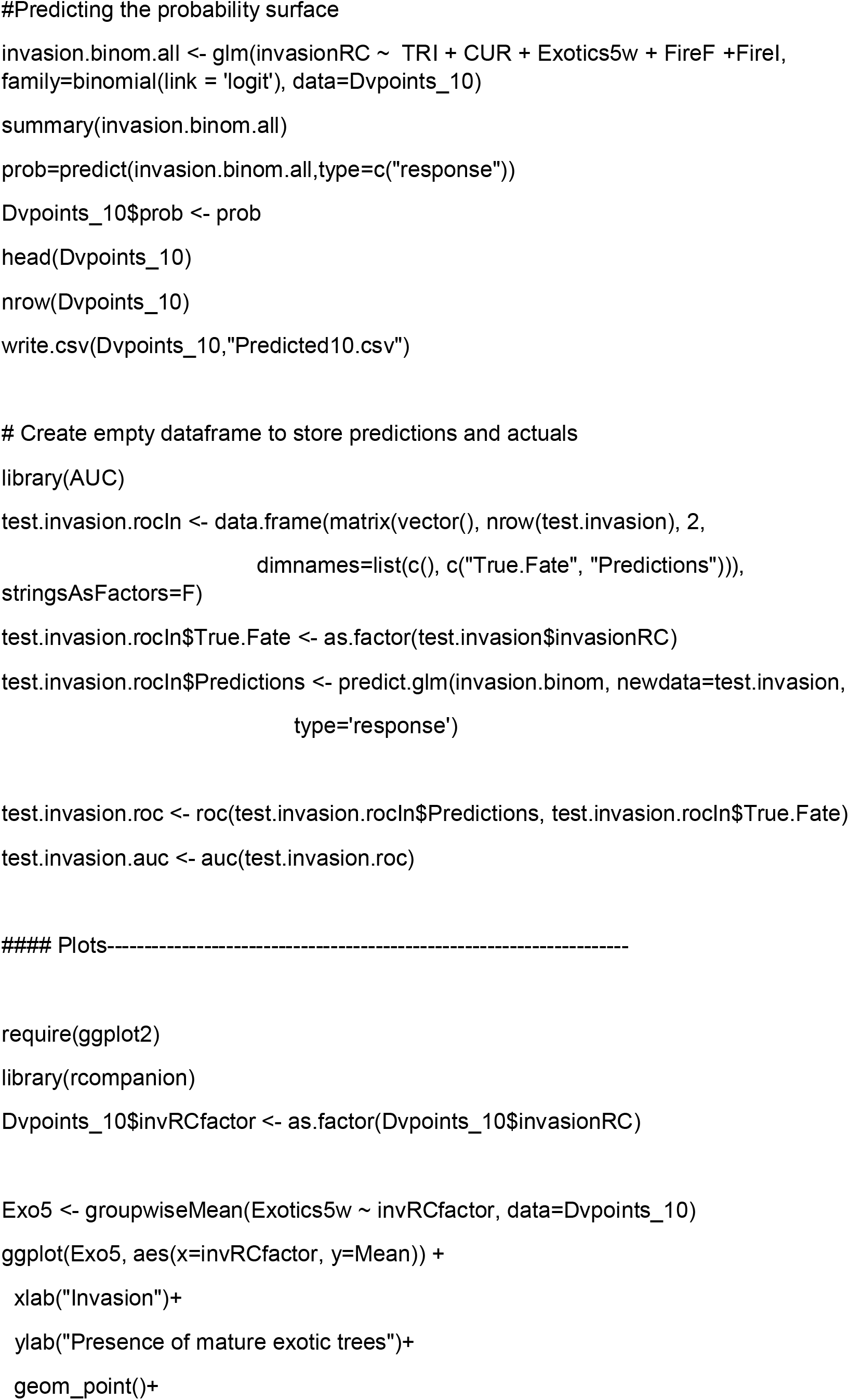
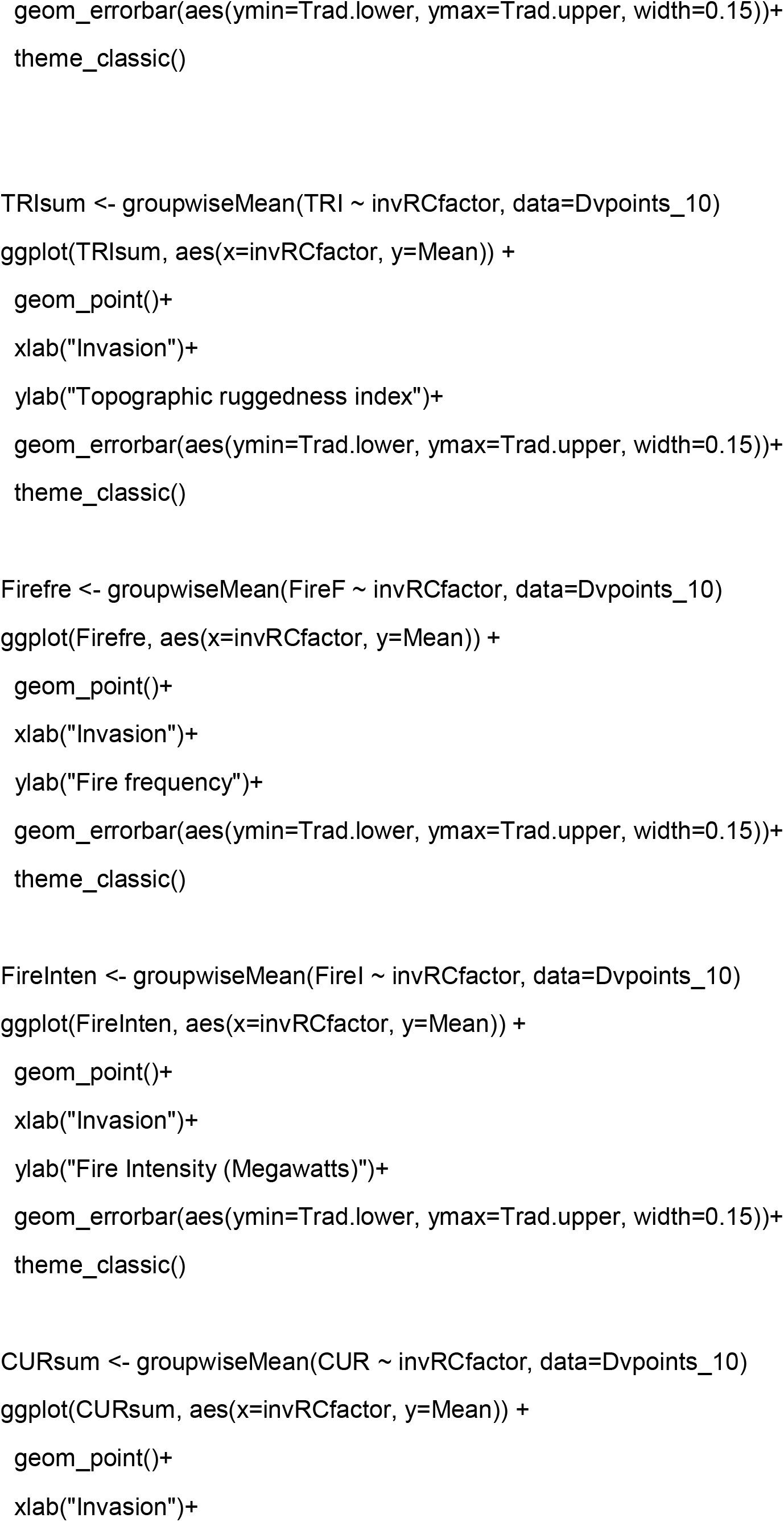
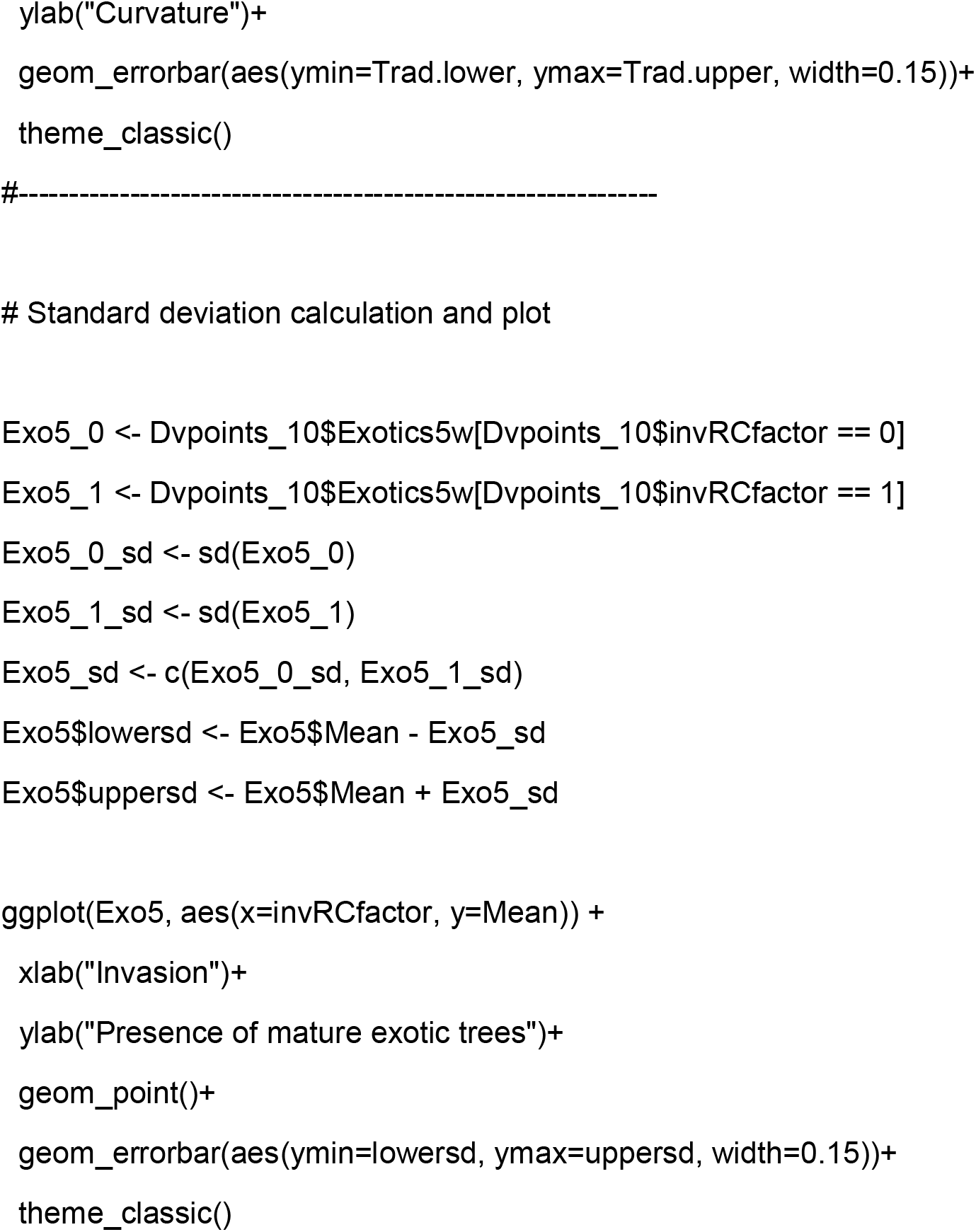

## Notes

### Competing Interest Statement

The authors have declared no competing interest.

